# Chronic paroxetine blunts stress response and normalizes adverse behavioral effects seen acutely in nulliparous rats

**DOI:** 10.1101/687509

**Authors:** Greta Tuckute, Agnete Overgaard, Vibe G. Frokjaer

## Abstract

Selective serotonin reuptake inhibitors (SSRI) are widely used antidepressants and their effect is partly mediated by restoring stress axis dynamics, which may depend on sex and hormonal states. In the present study, we investigate the effect of daily injections of the SSRI paroxetine (5 mg/kg s.c.) on swim stress-induced corticosterone (CORT) response, depressive-like behavior (forced swim test) and anxiety-like behavior (open field test) in nulliparous Sprague Dawley rats. Data were acquired after either 1-3 (acute PXT) or 11-13 (chronic PXT) injections. We found that chronic, but not acute, paroxetine blunted the swim stress-induced CORT response. We observed an increase in depressive-like and anxiety-like behavior following acute PXT, and a normalization after chronic PXT treatment. Intriguingly, our findings of rapid recovery from adverse SSRI effects differ from corresponding studies performed by our group in postpartum rats. Thus, the study emphasizes that mechanisms of action and efficacy of SSRIs differ according to reproductive states, which if translated to humans may inform treatment strategies, beyond SSRIs alone, for hormone transition related depressive states.

## 1. Introduction

During major depressive episodes, the hypothalamic-pituitary-adrenal (HPA) axis is usually hyperactivated with blunted dynamics (Ising et al., 2007; Ruhé et al., 2015). Selective serotonin reuptake inhibitors (SSRIs) decrease HPA-axis hyperactivity [1] and improve HPA system regulation by restoring HPA-axis negative feedback [2]. The efficacy of SSRIs on HPA dynamics has been linked to sex differences, with poorer response in hormone transition phase-specific periods including pregnancy and menopause [3]. Surprisingly, even though the lifetime risk for major depression is twice as high in women relative to men [4] most SSRI studies are performed in male animal models.

We have recently shown that treatment with the SSRI paroxetine (PXT) completely blunts the swim stress-induced stress response and induces depressive-like behavior in the forced swim test (FST) in a perinatal depression rat model [5], which points to an adverse effect of SSRIs when applied during the early postpartum period. However, we do not know whether SSRIs produce similar HPA axis response and adverse behavioral effects in nulliparae. Here, we determine the effect of PXT on corticosterone (CORT) response to swim stress, depressive-like behavior in the FST, and anxiogenic behavior in the open field test (OFT) in nulliparous rats.

## 2. Materials and methods

### 2.1 Animals

Eighteen female (250-275 g), age-matched Sprague-Dawley rats (Taconic, Denmark) were pair-housed under 12h:12h cycle (light on 0700) and given rat chow and tap water *ad libitum*. Rats were acclimatized for a week, assigned to treatment groups and handled for a week prior to the experimental start. Each cage belonged to the same experimental group. Protocols were in accordance with guidelines from The Danish Animal Ethics Council (license no. 2017-15-0201-01283).

### 2.2. Experimental design

Rats received daily injections (0900-1000) of either vehicle (5% dextrose in sterile water, 1 ml/kg s.c.) or the SSRI Paroxetine HCl (PXT; Carbosynth Limited; 5 mg/kg s.c.) for 13 days. PXT was brought into solution (5% dextrose in sterile water) using an ultrasonicator. Behavioral tests were performed at 1130-1500. Rats were randomly split into an acute PXT group (stress response measured on day 1), a chronic PXT group (stress response measured on day 11) and a vehicle group (stress response measured on day 11). Group size was n=6/group to obtain a power of 0.80, which was calculated based on the expected outcome of the stress response. Rats underwent two exposures in the forced swim test (FST), 24h apart, with the first exposure (FST1) used to measure the CORT response and the second exposure (FST2) given to examine behavior. The day after FST2, animals were tested in the open field test (OFT) (for experimental timeline, see Fig. 1A).

**Fig. 1.**
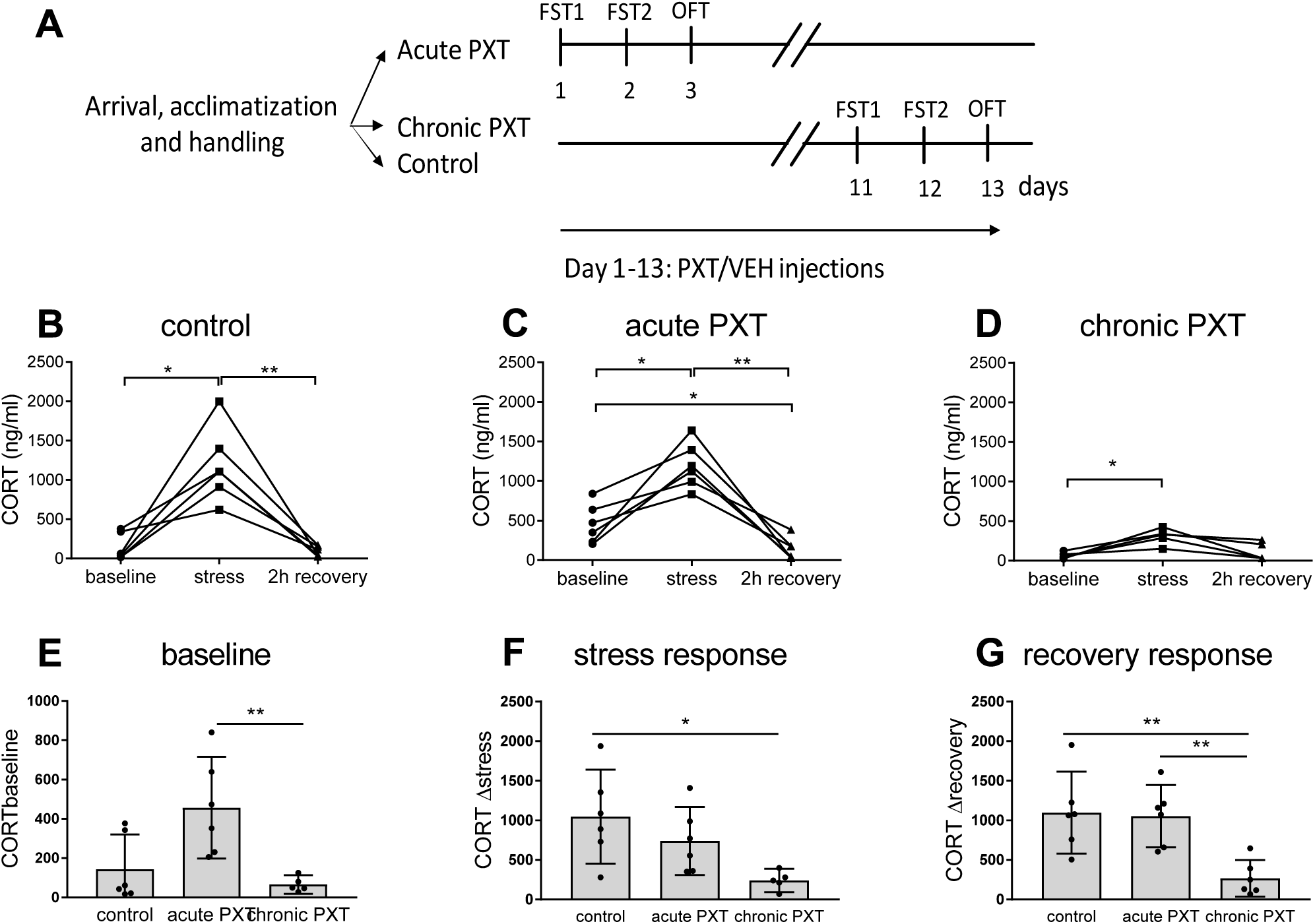
Experimental timeline and stress-induced corticosterone measurements. A) Timeline of experimental events. Rats were randomly assigned to one of three treatment groups: Acute PXT, chronic PXT and control. Rats were pair-housed and acclimatized for one week before injection start. The acute PXT group was tested on days 1-3, while the chronic PXT and control group were tested on days 11-13. Rats were under PXT treatment during testing. The acute PXT group received 1 PXT injection and the chronic PXT group received 11 daily s.c. PXT injections before the swim-stress induced CORT response measurement. B-D) Corticosterone at baseline, immediately after 15 min swim stress and after 2 h recovery. Graphs depict individual data points, connected for individual animals. All groups elicited a stress response, but only controls (B) and acute PXT (C) had significant recovery responses. Recovery CORT was below baseline level after acute PXT. E-G) Graphs depict mean with 95% CI and individual data points. E) CORT baseline measured 2.5-3 h after injection was increased after acute PXT compared to chronic PXT. F) Swim stress-induced CORT increase from baseline (Δ stress response) was reduced after chronic PXT relative to controls. G) CORT decrease from stress-induced CORT levels (Δ recovery response) was decreased after chronic PXT compared to controls. Results from Dunnett’s’s multiple comparisons test are marked with * for *p*<0.05, and ** for *p*<0.01. CORT, corticosterone; PXT, paroxetine; VEH, vehicle; FST, forced swim test; OFT open field test

### 2.3. Forced swim test (FST)

The FST was used to assess depressive-like behavior. A glass cylindrical tank (50 x 20 cm) was filled with water (25±0.1°C) to a depth of 35 cm, and light was dimmed during the test. FST1 was a 15min test performed at 1130-1500, and was used as an acute stressor. FST2, a 5min test the day after, assessed the behavioral response. Video recordings were manually scored for time spent immobile, swimming and climbing, by an observer unaware of treatment conditions.

### 2.4. Swim stress-induced HPA reactivity

Basal CORT levels were measured at 1130-1230, stress CORT levels were measured immediately after FST1, and recovery CORT levels were measured after 2h recovery in the home cage. Blood (max 100µl) was collected via tail vein puncture within 3 min of moving the cage or immediately following removal from the FST.

### 2.5. Open field test (OFT)

The OFT was used to assess locomotor activity and anxiety-like behavior. The 80×80 cm^2^ arena with 30 cm high walls was placed in a dimly lit room with a video camera installed above. Rats were placed in the arena facing a corner and activity was video recorded for 10 min. The videos were analyzed with EthoVision XT (Noldus Information Technology) to assess cumulative duration in center, center frequency, total distance travelled and time mobile. For the analysis, the arena was divided into 16 equal-sized squares, and the 4 central squares was defined as the center.

### 2.6. Corticosterone radioimmunoassay

Baseline, stress and recovery blood samples for the swim stress-induced CORT measurements were obtained from tail vein puncture. Blood was stored overnight at 4°C to allow blood to clot completely. Following centrifugation at 10000 *g* for 15 min, serum was collected and stored at −20°C. CORT was measured using ImmuChem Double Antibody 125I Radioimmunoassay Kit (MP Biomedicals). The cross-reactivity with other steroids is less than 0.4%. All samples were run in triplicates. The average intraassay coefficient of variation (CV) was 4.7%.

### 2.7. Data analyses

To determine whether a swim stress-induced CORT response was elicited, serum CORT was analyzed separately for each group, using repeated measures ANOVA with time (baseline, stress, recovery) as a within-subject factor. To test whether CORT baseline level, swim stress-induced CORT increase from baseline (Δ stress response), CORT decrease from stress-induced CORT levels (Δ recovery response), and FST and OFT measures differed between groups, we used a one-way ANOVA followed by a Dunnett’s multiple comparisons test, comparing chronic PXT with the two other groups. *P*<0.05 was considered significant.

### 2.8. Methodological considerations

Because the experimental design includes a vehicle group receiving chronic injections across 11 days and not a vehicle group for direct comparison with the acute PXT group, we restrict our analyses to chronic PXT vs acute PXT and chronic PXT to chronic vehicle groups. Thus, we interpret differences between acute PXT and vehicle groups with longer injection exposure time with caution.

## 3. Results

### 3.1. Chronic paroxetine blunted swim stress-induced corticosterone response

The stress-induced CORT response was measured during the first exposure to FST. As expected, the control group elicited a significant stress response and full CORT recovery post stress (Fig. 1B). The acute PXT group showed an increased CORT baseline (Fig. 1E) compared to the chronic PXT group (*p*<0.01), and the acute PXT group also recovered to CORT levels below baseline (Fig. 1C). Chronic PXT treatment resulted in a highly blunted swim stress-induced CORT response, and reduced CORT recovery levels post stress relative to both control and acute PXT (*p*<0.01; Fig. 1G).

### 3.2. Acute paroxetine increased anxiety-like behavior in the open field test and increased depressive-like behavior in the forced swim test

In the OFT, percent time spent in center (Fig. 2A) and center frequency (Fig. 2A) was significantly decreased by acute PXT treatment compared to chronic PXT treatment (*p*<0.05), whereas distance travelled in the OFT was not affected (*p*>0.99; Fig. 2A).

**Fig. 2.**
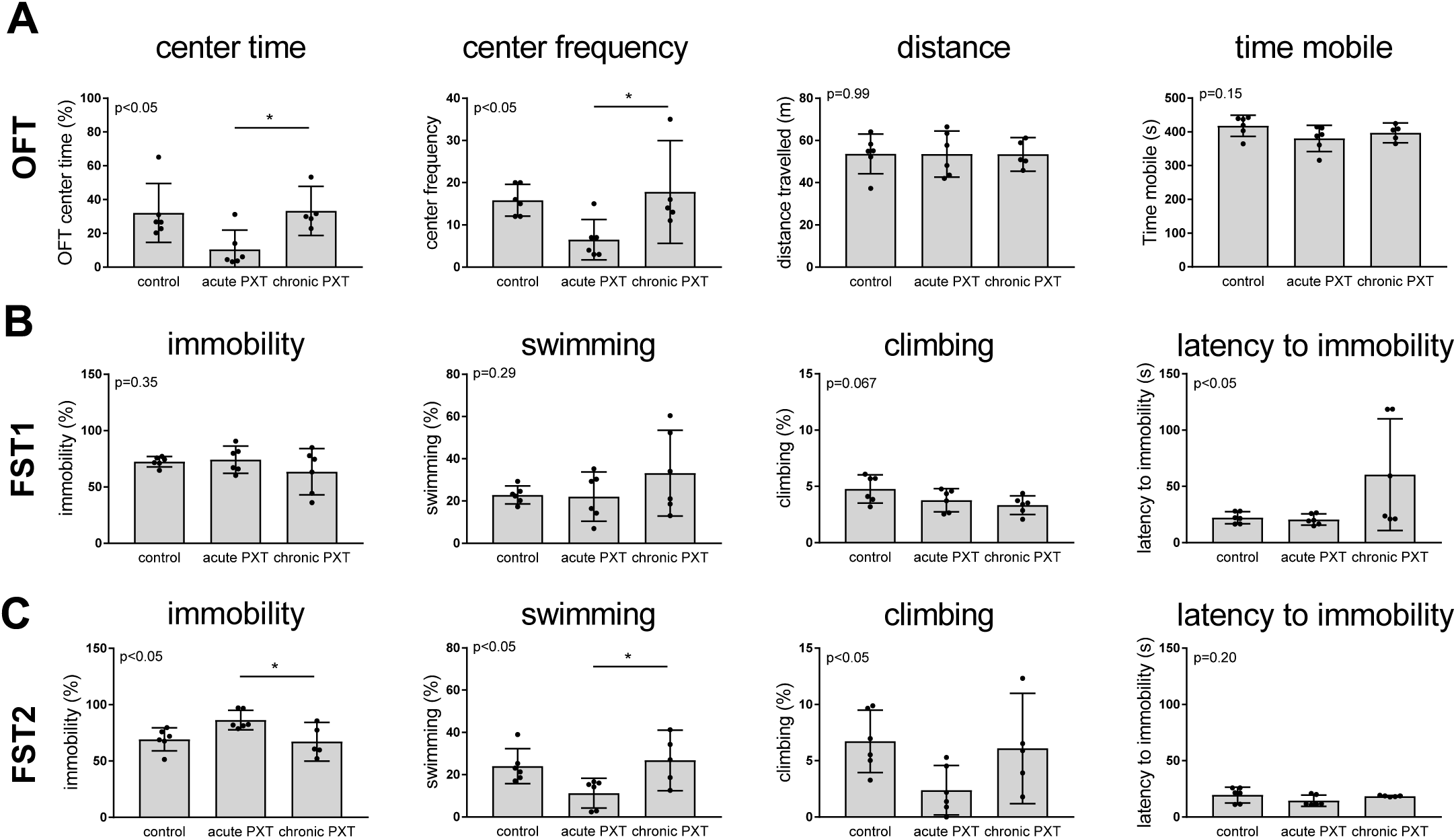
Anxiety- and depressive-like behavior. A) Open field test (OFT). Percent time spent in the center in the OFT and OFT center time was decreased by acute PXT compared to chronic PXT. Distance travelled and time spent mobile was not affected by PXT. B) Forced swim test 1 (FST1). There was an overall effect on latency to immobility and an overall trend in percent time spent climbing, with chronic PXT tending to decrease climbing but increase latency to immobility. C) FST2. Acute PXT increased depressive-like behavior as shown by increased immobility and decreased swimming and climbing relative to chronic PXT. Chronic PXT did not significantly alter behavior compared to the control group. Results from Dunnett’s multiple comparisons test are marked with * for *p*<0.05. Graphs depict mean with 95% CI and individual data points. PXT = paroxetine; VEH = vehicle; OFT = open field test; FST = forced swim test.

In FST2, acute PXT treatment significantly increased percent time spent immobile and decreased time spent swimming compared to chronic PXT treatment (*p*<0.05) and affected climbing (Fig. 2C; *p*<0.05). These parameters were normalized after chronic PXT treatment, and no significant differences between the control group and chronic PXT group were found (Fig. 2C).

In FST1, chronic PXT treatment increased latency to immobility (*p*<0.05) and tended to decrease climbing (*p*=0.067; Fig. 2C).

## 4. Discussion

We demonstrate that chronic, but not acute, PXT blunts the swim stress-induced stress response in nulliparous rats. Moreover, we find an increase in depressive- and anxiety-like behavior after acute, but not chronic, PXT administration.

### Chronic paroxetine reduced the stress-induced corticosterone response

We demonstrate that acute PXT does not affect the ability to mount a swim stress-induced response in nulliparous rats, whereas the stress response is blunted by chronic PXT. In the acute PXT group, we noted an increase in baseline CORT levels and below baseline CORT recovery levels relative to a suboptimal reference group, which may suggest that an acute PXT injection increases CORT, and that CORT levels are not completely back to baseline levels 3 hours following the first PXT injection. This effect is not observed after chronic PXT, supporting the notion that SSRIs dampen HPA system tone only after chronic administration [6]. While we cannot exclude that the repeated injections *per se* decrease CORT baseline, this finding is consistent with the adverse behavioral effects of acute PXT we observe in FST and OFT.

In the present study, we observe a higher stress response in nulliparous rats compared to earlier observations in postpartum rats in similar experimental designs [5], which parallels to earlier studies showing a dampened HPA-axis during the postpartum period in both rats [7] and women [8]. Previously we found that the CORT swim stress-induced response is completely blunted in postpartum rats by both acute *and* chronic PXT [5]. In this study, we demonstrate a small but sustained significant stress response after chronic PXT, which indicates that blunting of the CORT stress response after PXT administration is less profound in nulliparous rats compared to our previous postpartum observations [5]. This finding might be related to altered CORT dynamics postpartum. Thus, the chronic effect of SSRIs might be beneficial in terms of ameliorating depressive-like behavior in nulliparous, but not so in postpartum rats. Our finding of an effect of chronic PXT on HPA-axis function is consistent with clinical findings demonstrating an improvement of HPA system function after 2-3 weeks of SSRI administration in depressed patients, which appear to be critical to the antidepressant effects of SSRIs [1]. Moreover, the different effect of PXT on HPA-axis dynamics in nulliparous and postpartum rats demonstrates that findings from nulliparous rat models of depression do not translate to the postpartum, and that mechanisms of SSRIs on the postpartum brain must be investigated separately.

### Acute paroxetine increased depressive-like and anxiety-like behavior

We demonstrated that acute PXT treatment increased depressive- and anxiety-like behavior in nulliparous rats relative to chronic PXT treatment.

In male rat models of depression, SSRIs reduce immobility in the FST after chronic treatment at the same low dose as used in our study [9]. In female rats, however, chronic SSRI administration fail to affect immobility in the FST in either nulliparous, postpartum or ovariectomized rats [10]–[13]. In line with these studies, we observed no effect of chronic PXT on depressive-like behavior in nulliparous rats. In postpartum rats, SSRIs have recently been demonstrated to increase depressive-like behavior [5], [14]. Thus, the effects of SSRIs on depressive-like behavior differ with sex and reproductive state in rat models. Our finding of an initial triggering of depressive-like behavior due to PXT treatment in nulliparae is contrasted by a recent study demonstrating that depressive-like behavior is still increased after 13 daily PXT injections in the postpartum period [5]. However, studies of the late postpartum show no difference in FST-related immobility between SSRI treated (fluoxetine) and control dams [13]. Thus, our data suggest that the adverse reaction period with triggering of depressive-like behavior due to SSRI exposure is shorter in nulliparous rats compared to postpartum rats. Taken together, we propose that SSRIs have distinct effects and reduced efficacy on the postpartum (depressed) brain, which may be related to dampening of HPA dynamics.

In the OFT, acute, relative to chronic PXT administration increased anxiety-like behavior, which indirectly aligns with clinical data suggesting that patients experience an initial increase in anxiety when given SSRIs [6]. Likewise, this finding is consistent with studies demonstrating anxiogenic effects of acute, but not chronic, SSRIs in male rats [15]. In contrast, SSRIs induce anxiogenic behavior in postpartum rats across the early to mid-postpartum (Overgaard et al., unpublished observation), as well as during the late postpartum after chronic administration [12], [14]. Chronic SSRIs in nulliparous rats do not induce anxiety-like behavior [11], [16]. Taken together, this suggests that male and female rats have similar anxiogenic response to SSRIs, except for females in the postpartum, where the response seems to be delayed and prolonged.

In conclusion, we show that depressive- and anxiety-like behavior is increased after acute PXT relative to chronic PXT in nulliparous rats, which we propose is related to altered HPA axis dynamics after PXT administration. These data contrast earlier findings of ours from similar studies in postpartum rats, and underscore that the action of SSRIs is tightly linked to sex and reproductive status.

Thus, future studies must address risk and treatment mechanism specific to the postpartum brain, to provide improved clinical care for and prevention of perinatal depression.

